# Holotomography-driven learning for in-silico staining of single cells in flow cytometry avoiding co-registration

**DOI:** 10.1101/2025.07.22.666145

**Authors:** Daniele Pirone, Giusy Giugliano, Michela Schiavo, Annalaura Montella, Martina Mugnano, Vincenza Cerbone, Maddalena Raia, Giulia Scalia, Ivana Kurelac, Diego Luis Medina, Lisa Miccio, Mario Capasso, Achille Iolascon, Pasquale Memmolo, Pietro Ferraro

**Affiliations:** CNR-ISASI, Institute of Applied Sciences and Intelligent Systems “E. Caianiello”, Via Campi Flegrei 34, 80078 Pozzuoli, Napoli, Italy; Department of Mathematics and Physics, University of Campania “Luigi Vanvitelli”, Viale Abramo Lincoln 5, 81100 Caserta, Italy; TIGEM, Telethon Institute of Genetics and Medicine, Via Campi Flegrei 34, 80078 Pozzuoli, Napoli, Italy; Department of Advanced Biomedical Science, University of Naples “Federico II”, Via Sergio Pansini 5, 80131 Napoli, Italy; CEINGE - Advanced Biotechnologies, Via Gaetano Salvatore 486, 80131 Napoli, Italy; DMMBM, Department of Molecular Medicine and Medical Biotechnology, University of Naples “Federico II”, Via Pansini 5, 80131 Napoli, Italy; DICMaPI, Department of Chemical, Materials and Production Engineering, University of Naples “Federico II”, Piazzale Tecchio 80, 80125 Napoli, Italy; DIMEC, Department of Medical and Surgical Sciences, Alma Mater Studiorum-University of Bologna, 40138 Bologna, Italy; IRCCS Azienda Ospedaliero-Universitaria di Bologna, 40138 Bologna, Italy; Medical Genetics Unit, Department of Medical and Translational Science, University of Naples “Federico II”, Via Sergio Pansini 5, 80131 Napoli, Italy

## Abstract

Virtual staining is the current state-of-the-art computational technique to cleverly enhance intracellular specificity in unstained biological samples by using convolutional neural networks (CNNs) trained on co-registered pairs of unstained/stained images. While effective, this approach suffers from unpredictable biases inherent to fluorescence microscopy and encounters challenges when applied to flow cytometry data as it would require accurate co-registration on a huge number of images. Here, we present a novel method that exploits for the first time a Holotomography-driven learning to completely eliminate the need for co-registration. We demonstrate that training a CNN on a stain-free dataset of 3D refractive index tomograms of flowing cells elegantly unlocks stain-free intracellular specificity in quantitative phase imaging flow cytometry. This breakthrough, by circumventing the critical co-registration bottleneck, opens unprecedented perspectives for label-free, high-throughput imaging flow cytometry, offering a powerful new paradigm for advanced 2D and 3D single-cell analysis.

## Introduction

Cellular populations are often heterogeneous in terms of size, shape, physiological state, and cell cycle phase [1]. Common measurement devices typically provide only global parameters about the entire cellular population, while failing to capture the cell-to-cell differences and the individual behaviors that instead could be crucial for understanding a biological phenomenon [2]. Thus, advanced single-cell analysis is considered the new frontier in omics enabling to transform systems biology [3]. For many decades, Fluorescence Microscopy (FM) has been the gold standard technique for imaging and measuring single-cell properties through the employment of various stains able to tag intracellular organelles and biochemical processes [4], especially with the advent of super-resolution strategies [5]. However, FM has some intrinsic issues that can perturb the single-cell imaging and its downstream analysis, such as photobleaching, phototoxicity and high signal-to-noise ratio (e.g., due to autofluorescence or spectral overlap between multiple fluorophores), and is often affected by sample preparation- and operator-dependance [6].

For this reason, more recently, label-free microscopes [7] have emerged as a promising alternative to FM for the unperturbed analysis of biological samples in life science research [8]. In particular, Quantitative Phase Imaging (QPI) has been gaining momentum [9], with a remarkable acceleration in the recent years, also thanks to the combination with Artificial Intelligence (AI) [10]. A quantitative phase map (QPM) relies on endogenous image contrast based on the optical phase delay introduced by the biological sample on an incident wavefront, which allows to avoid any exogenous staining. Moreover, QPI gives access to a quantitative biophysical measurement of the sample as its morphology and its refractive index (RI) distribution are mapped together in the same 2D image [9]. Therefore, QPI-based applications in biology and biomedicine have rapidly multiplied and improved over time [9,11]. In addition, the latest advancement of QPI in 3D — commonly known as Tomographic Phase Microscopy or Holographic Tomography (HT) — offers the unique capability to reconstruct the volumetric RI distribution of biological specimens by combining multiple QPMs acquired from various viewing angles [12–15], reaching the nanometer scale [16]. Among the several HT implementations [13–15], the imaging flow cytometry demonstration represents the most promising strategy to simultaneously achieve high-throughput, 3D and stain-free analysis at the single-cell scale [17,18]. In Holo-Tomographic Flow Cytometry (HTFC), the 2D QPMs of single cells flowing and rotating along a microfluidic channel are acquired at multiple viewing angles, thus enabling the reconstruction of 3D RI tomograms [17]. Several applications based on HTFC have been demonstrated [19–22], including a proof-of-concept of label-free liquid biopsy [23].

Despite the remarkable advancements made in QPI, lack of intracellular specificity remains its main limiting factor in respect to FM. In fact, in the absence of specific fluorescence stain, phase-contrast imaging is the only guide to recognize specific intracellular compartments. However, this is a non-trivial process due to the high similarities among the RI distributions of different organelles [24]. Therefore, several in-silico staining strategies have been developed in recent years to bridge the gap with FM in terms of molecular specificity [25–27]. Among the others, the virtual staining concept has proven to be a very clever and effective solution [28]. Virtual staining relies on deep learning in a way that Convolutional Neural Network (CNN) learns to emulate the fluorescence signal in stain-free images after being trained with a dataset of paired co-registered unstained/stained images. While the virtual staining paradigm has found applications in many label-free microscopes [28,29], in the specific field of QPI it has been applied to retrieve specificity in the 2D QPMs of unlabeled tissues [30], as well as from 2D QPMs [31,32] and 3D RI tomograms [33,34] of unlabeled single cells, culminating in the development of a label-free artificial confocal microscope [35].

Despite virtual staining has marked a turning point in the development of new QPI-based applications, training a CNN on an FM dataset has not achieved full independence of label-free QPI from FM imaging and its inherent limitations. In fact, alongside molecular specificity, the CNN also learns FM-related artifacts. Moreover, acquiring a dataset of co-registered QPI/FM images is particularly challenging in flow cytometry settings. For this reason, a completely different in-silico staining approach has recently been introduced for organelle segmentation, termed Computational Segmentation based on Statistical Inference (CSSI) [36–38]. The CSSI algorithm was designed to recognize statistical similarities among intracellular RI values inherently mapped in a stain-free phase-contrast tomogram, thus avoiding any training with FM data. However, the CSSI algorithm has very heavy computational burdens and it cannot be implemented in 2D QPMs because the lack of a sufficient number of pixels prevents reliable statistical similarity testing, which is instead the core of the CSSI [36–38].

To address the main limitations of virtual staining and the CSSI algorithm, here we introduce a novel in-silico staining approach based on an innovative Holotomography-driven learning framework. Notably, this framework is, for the first time, trained on a fully stain-free dataset of 3D RI tomograms of flowing cells, segmented by CSSI in a self-supervised learning setting [39–42], thus avoiding any experimental co-registration from fluorescence maps (Fig. 1). Using nucleus segmentation as a test case, we demonstrate that the proposed method achieves two significant objectives: (1) it enables accurate real-time retrieval of nuclear specificity in 2D QPMs of single cells analyzed in a flow cytometry setting; (2) it drastically reduces the computational cost in HTFC compared to CSSI, cutting computational time by up to four orders of magnitude for segmenting nuclei in 3D RI tomograms — from tens of minutes to fractions of a second. Importantly, with respect to objective (1), it is worth highlighting that intracellular specificity has not yet been achieved in 2D QPI flow cytometry (QPIFC) to date. For training and testing the Holotomography-driven CNN, by using the HTFC experimental system, we collected a dataset of 23,399 QPMs corresponding to 220 cervical cancer cells (HeLa) and 301 breast cancer cells (MCF-7). In particular, we employ the dataset of HeLa cells to train and test the CNN, and then we demonstrate that, remarkably, the same trained Holotomography-driven architecture is able to generalize in-silico staining of nucleus in the MCF-7 cell line never seen during training. We publicly share the stain-free QPM dataset used in this study, thus providing for the first time an openly available dataset of QPMs acquired in flow cytometry mode. Finally, in addition to the conventional CNN assessment, we compare our results with the 2D FM measurements obtained by a commercial Fluorescence Imaging Flow Cytometry (FIFC) system, thus demonstrating the robustness of our method.

**Fig. 1.**
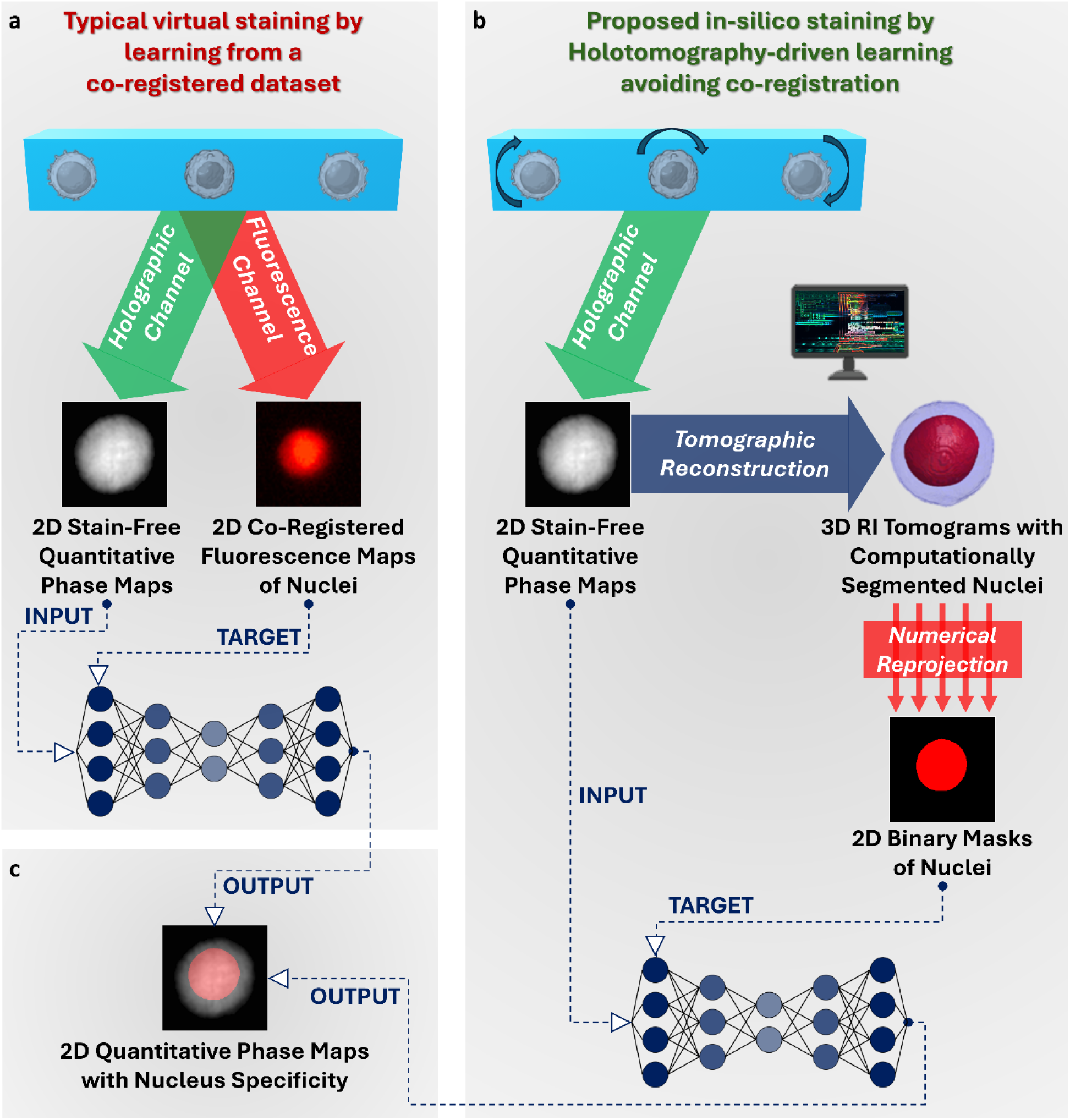
Conceptual comparison between the conventional virtual staining and the proposed Holotomography-driven staining. **a** Virtual staining is based on experimental co-registration between QPMs and fluorescence maps to create the dataset for CNN training. **b** Holotomography-driven staining is based on HTFC to create a stain-free dataset for CNN training, thus employing only QPMs and avoiding any experimental co-registration with fluorescence maps. **c** Both methods allow to retrieve intracellular specificity in stain-free QPMs of single cells in flow cytometry.

We believe that the proposed Holotomography-driven staining could represent a breakthrough not only for QPIFC and HTFC, but for all label-free fields in which FM still represents an indirect ground truth, thus allowing to definitively break these ties and move towards a purely label-free microscopy.

## Results

### Stain-free training of the Holotomography-driven CNN for the in-silico staining of 2D QPMs of flowing cells

A scheme of the Holotomography-driven learning for in-silico staining is sketched in Fig. 2 and illustrated in Supplementary Video 1. The HTFC system described in the Methods section and the corresponding numerical processing are employed to collect the 2D QPMs of single cells flowing and rotating in suspension along a microfluidic channel. Thanks to the HTFC concept, a stack of QPMs is available for each cell containing different views because of its roto-translation during the holographic acquisition. For this reason, the 3D RI tomograms can be reconstructed at the single-cell level. Then, the implementation of the CSSI algorithm allows to segment the 3D volume of stain-free nuclei in suspended cells [36]. The CSSI algorithm is indeed proved to retrieve the nucleus specificity by avoiding the external FM guide, while exploiting the statistical inference among RI values to cluster intracellular voxels into specific organelles [36–38]. Then, the 3D nuclear volumes segmented by the CSSI algorithm are reprojected along the same viewing angles at which the 2D QPMs have been experimentally imaged. In this way, a fully stain-free dataset of 2D nucleus masks is created, corresponding to the experimental QPMs. Such QPMs are finally used as input of a CNN aimed at segmenting the 2D nucleus regions after having been trained through the reprojected 2D nucleus masks used as target.

**Fig. 2.**
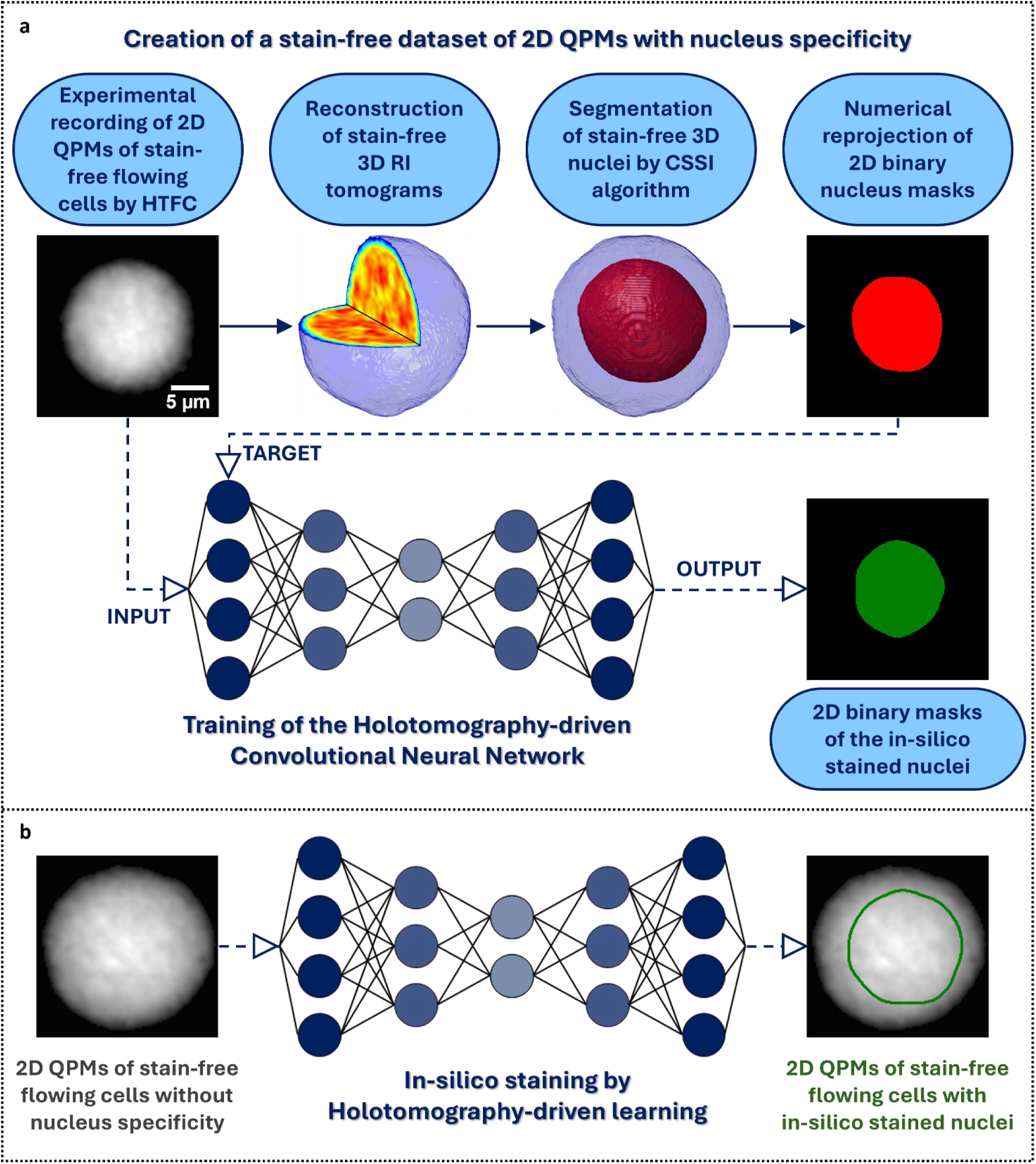
Workflow of the Holotomography-driven learning for in-silico staining (Supplementary Video 1). **a** A stain-free dataset of 2D QPMs of flowing cells with nucleus specificity is created by combining the HTFC experiments and the CSSI algorithm, at the aim of training a Holotomography-driven CNN. **b** Starting from the input stain-free QPM, the Holotomography-driven CNN allows to predict the 2D nucleus mask.

For training the Holotomography-driven CNN, a dataset of 220 HeLa cells was collected (Table S1). To avoid redundancies, only QPMs limited to a range of 360° of rotation angles (i.e. viewing angles) were selected. In this way, 10,729 QPMs were considered, i.e. around 49 QPMs per cell.

It is worth noting that the cellular roto-translation at the basis of HTFC not only enabled the creation of a 2D stain-free dataset, but also allowed to experimentally augment the number of observations as, for each cell, tens of viewing angles were imaged. The stain-free dataset of segmented QPMs was then divided into training set, validation set, and internal test set, according to the numbers reported in Table S1. A CNN model derived from a DeepLabv3+ ResNet-18 architecture [43] was trained in order to validate the proposed strategy (as reported in Methods section). The CNN model took in input the QPM and provided in output a binary mask in which only the nucleus area is set to 1, as shown in Fig. 3a-d (first three columns). Then, after predicting the 2D nucleus mask, its contour was overlapped to the input QPM in order to enable downstream quantitative analysis at the single intracellular level (see last column in Fig. 3a-d).

**Fig. 3.**
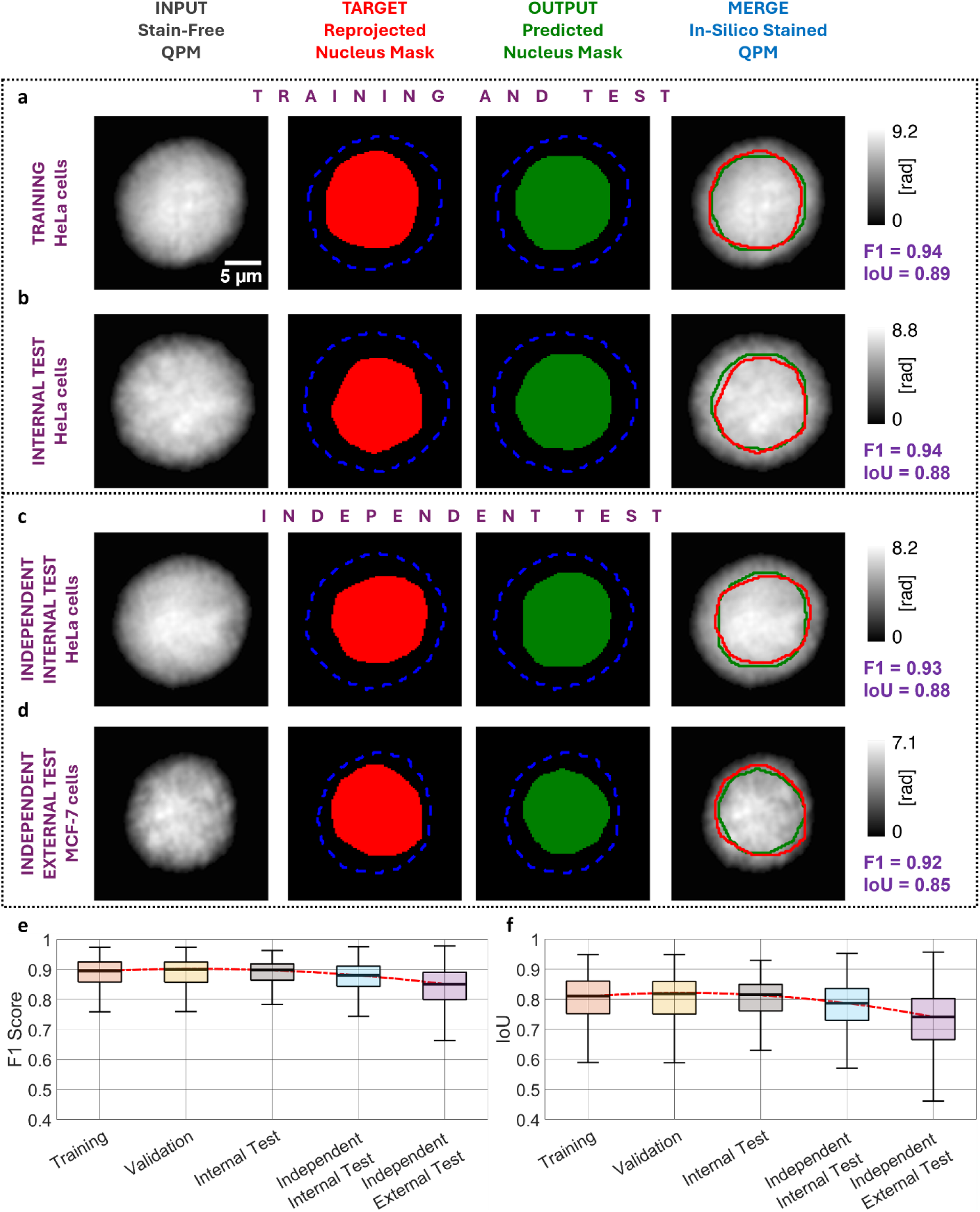
Holotomography-driven staining of the 2D QPMs of stain-free flowing cells. **a-d** Examples extracted from the training set, internal test set, independent internal test set, and independent external test set, respectively. From left to right, input QPM recorded by HTFC, target nucleus mask (red pixels are 1 and blue dashed line is the cell contour) reprojected after CSSI, output nucleus mask (green pixels are 1 and blue dashed line is the cell contour) predicted by CNN, and in-silico stained QPM with overlapping the contours of the target (red) and output (green) nucleus masks. The F1 scores and the IoU values are reported on the right. **e,f** Boxplots of the F1 scores and IoU values about the training, validation, internal test, independent internal test, and independent external test sets. For each boxplot, the central line is the median, the bottom and top edges of the box are the 25th and 75th percentiles, respectively, and the whiskers extend to the most extreme non-outlier points. The red dotted line is a quadratic fitting of the median values. Images (a-d) are approximately the 90th percentile of the IoU distributions for their datasets.

In Fig. 3a,b, we show an example of CNN prediction on two HeLa cells belonging to the training set and the internal test set, respectively. It can be seen that the CNN was able to predict a nucleus contour consistent with the target one. To quantitatively evaluate this, we measured two performance metrics described in Methods section, i.e. the F1 score and the Intersection over Union (IoU). The F1 score is the harmonic mean between recall and precision, and is very useful with unbalanced classes, such as the segmentation problem herein addressed (the background area is much greater than the nucleus area). As reported in the boxplots of Fig. 3e, the trained CNN model reached F1=0.89±0.04 over the 7,297 QPMs of the training set, F1=0.89±0.05 over the 2,418 QPMs of the validation set, and F1=0.88±0.05 over the 1,014 QPMs of the internal test set. Instead, the IoU is a very common metric to assess performance of a segmentation network, as it measures the ratio of the intersection and union areas between the target and predicted nucleus masks. The IoU metric is very sensitive to small discrepancies between target and prediction. Thus, even small differences can cause a significant reduction in the IoU value, making it difficult to reach very high values, especially in complex segmentation scenarios. To further confirm this, in Fig. S1, we report that small translations of a 2D object cause big reductions in the IoU values. Usually, IoU>0.5 is considered a good score for segmentation problems, and in particular 0.7<IoU<0.95 is a high score and IoU>0.95 is an excellent score. As reported in Fig. 3f, our Holotomography-driven CNN reached high values of IoU over the training set, validation set, and internal test set, i.e. IoU=0.80±0.07, IoU=0.80±0.07, and IoU=0.80±0.08, respectively.

Furthermore, we performed a robust assessment of the generalization ability of the proposed Holotomography-driven staining in two different ways. First, we assessed the internal generalization by performing another experiment on the HeLa cell line (Fig. 3c), completely independent of the HeLa experiment initially used to train, validate, and test the CNN, thus collecting data from 103 new HeLa cells, corresponding to 4,415 new QPMs (Table S1). In this independent internal test, we obtained F1=0.87±0.05 (Fig. 3e) and IoU=0.78±0.08 (Fig. 3f), consistent with the internal test performance. Finally, to assess the external generalization, we next carried out an additional experiment with a cell line never seen during training, i.e. the breast cancer MCF-7 (Fig. 3d), by acquiring 198 cells and the corresponding 8,255 QPMs (Table S1). Surprisingly, performances of this independent external test were in line with the other ones, i.e. F1=0.84±0.07 (Fig. 3e) and IoU=0.73±0.10 (Fig. 3f). The excellent generalization feature was also confirmed by the quadratic fitting overlapped to the boxplots in Fig. 3e,f, which shows a very weak decrease across the training, validation, internal test, independent internal test, and independent external test sets.

### Extension of the Holotomography-driven staining to the 3D RI tomograms of stain-free flowing cells

In the previous section, we have shown that the Holotomography-driven learning allows to predict the stain-free nucleus masks in the 2D QPMs of single cells flowing in suspension along a microfluidic channel. However, the same strategy can be also employed to segment the nucleus mask in their 3D RI tomograms, by following the pipeline described in Fig. 4 and illustrated in Supplementary Video 1. In fact, the HTFC paradigm, described in the Methods section, allows to reconstruct the 3D RI tomograms of single cells in flow cytometry environment if their rotation is additionally induced and imaged [19–23]. Currently, to the best of our knowledge, the sole fully computational method to retrieve the nucleus from the 3D RI tomograms of flowing cell is the CSSI algorithm [36], herein employed to create the stain-free dataset for training the in-silico staining of 2D QPMs. However, while CSSI is an intrinsically stain-free solution, the high computational burden limits its applications in flow cytometry, where a high throughput analysis of large cell numbers and respective big datasets is required. Indeed, to segment one single nucleus, the CSSI method takes tens of minutes on the 3D arrays herein considered. Moreover, the computational time increases in a nonlinear way with the spatial resolution of the 3D reconstruction, as well as its RAM memory requirements. A solution to this issue can be provided by Holotomography-driven staining by first predicting the 2D nucleus masks in the entire sequence of QPMs containing the cell’s roto-translation, and then by implementing the tomographic algorithm through the binary nucleus masks used as input. In this way, we obtain the 3D binary mask of the nucleus, which can be overlapped to the 3D RI tomogram in order to perform a quantitative measurement at the cellular and intracellular level. In our system, we employ the Filtered Back Projection (FBP) algorithm [44] to combine tens of 2D nucleus masks per cell. Therefore, the computational time of the FBP algorithm summed to the prediction time of the 2D nucleus masks (2.7 ms per QPM) results in a 3D nucleus segmentation performed in few tenths of a second instead of tens of minutes.

**Fig. 4.**
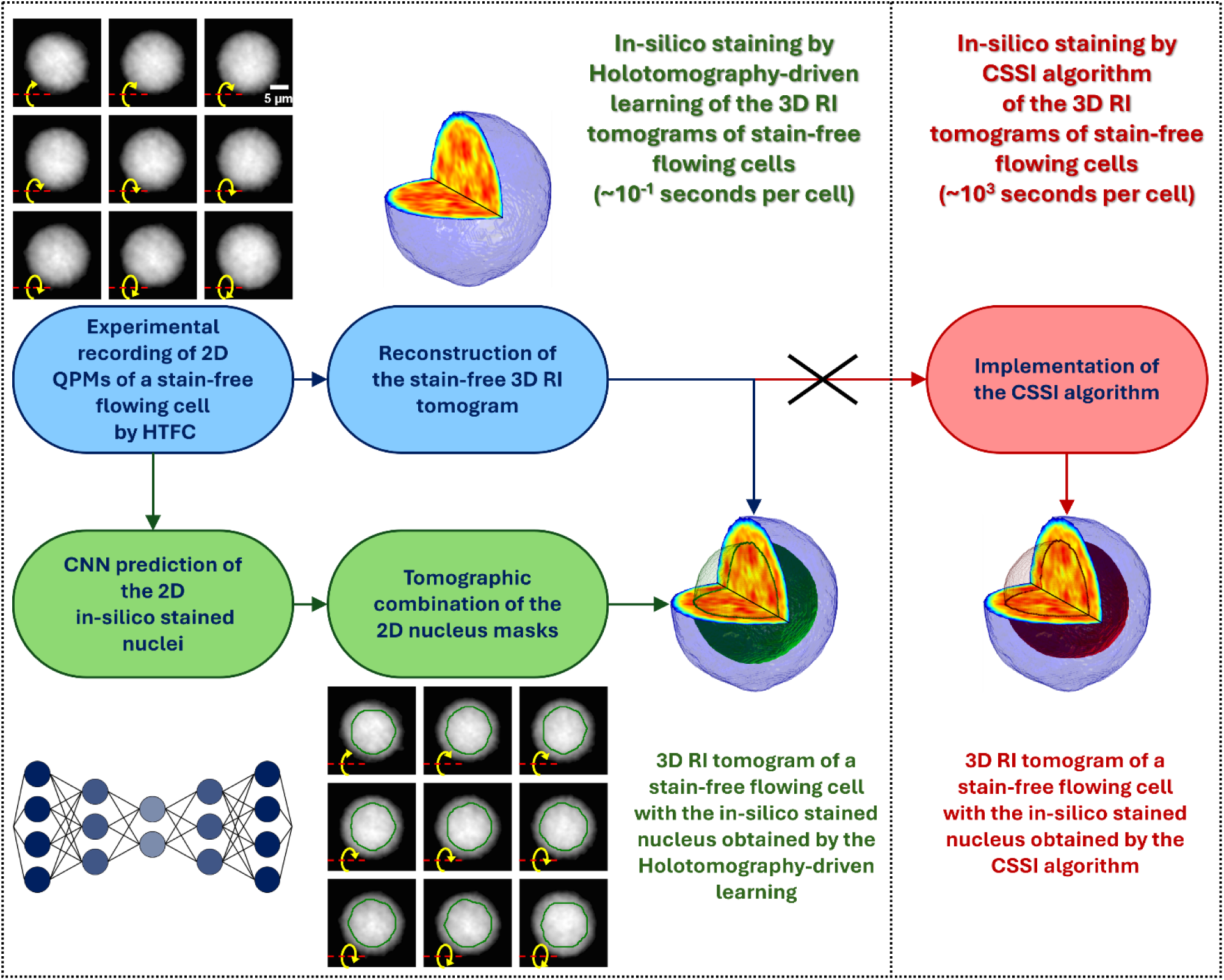
Scheme of the 3D extension of the Holotomography-driven staining (Supplementary Video 1). A stack of 2D QPMs containing the rotation of a stain-free flowing cell is given in input to the Holotomography-driven CNN, which provides the corresponding 2D nucleus masks in output. The stack of 2D nucleus masks is combined by a tomographic algorithm to compute the 3D nucleus mask of the flowing cell, which can be overlapped to its 3D RI tomogram computed in parallel from the same stack of 2D QPMs. This pipeline allows avoiding the implementation of the CSSI algorithm after the calculation of the 3D RI tomogram for segmenting its 3D nucleus mask, thus speeding up the 3D nucleus identification by four orders of magnitude.

In Fig. 5a-d, we show the CNN results of the 3D nucleus segmentation based on the Holotomography-driven learning. In the first column, the reconstructed 3D RI tomogram of the stain-free cell is reported, from which the 3D nucleus mask was segmented by the CSSI algorithm and used as ground-truth (second column). In the third column, the 3D nucleus mask obtained by combining the 2D in-silico staining predictions is reported, which was directly compared with its ground-truth in the last column, overlapped to the 3D RI distribution. Also in the 3D case, we considered the F1 score and the IoU as performance metrics of the in-silico staining segmentation.

**Fig. 5.**
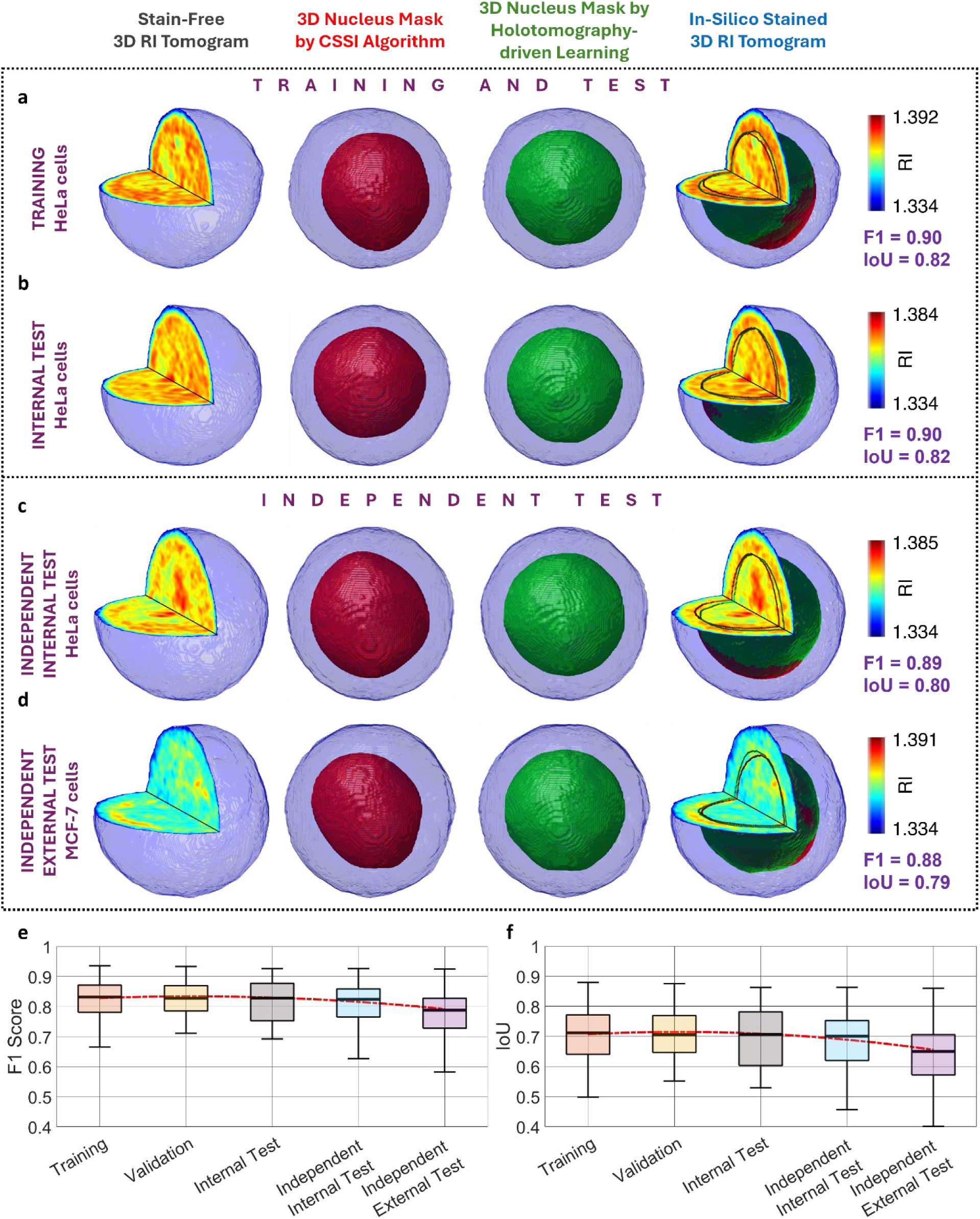
Holotomography-driven staining of the 3D RI tomograms of stain-free flowing cells. **a-d** Examples extracted from the training set, internal test set, independent internal test set, and independent external test set, respectively. From left to right, input RI tomogram recorded by HTFC, target nucleus mask (red) computed by CSSI within the cell shell (blue), output nucleus mask (green) computed by combining 2D CNN predictions within the cell shell (blue), and in-silico stained RI tomogram with overlapping the contours of the target (red) and output (green) nucleus masks. The F1 scores and the IoU values are reported on the right. **e,f** Boxplots of the F1 scores and IoU values about the training, validation, internal test, independent internal test, and independent external test sets. For each boxplot, the central line is the median, the bottom and top edges of the box are the 25th and 75th percentiles, respectively, and the whiskers extend to the most extreme non-outlier points. The red dotted line is a quadratic fitting of the median values. Images (a-d) are approximately the 90th percentile of the IoU distributions for their datasets.

In particular, as reported in the boxplots of Fig. 5e,f, respectively, we reached F1=0.82±0.06 and IoU=0.71±0.09 over the 150 HeLa tomograms of the training set, F1=0.82±0.06 and IoU=0.70±0.09 over the 50 HeLa tomograms of the validation set, F1=0.82±0.07 and IoU=0.70±0.11 over the 20 HeLa tomograms of the internal test set, F1=0.81±0.07 and IoU=0.68±0.09 over the 103 HeLa tomograms of the independent internal test set, and F1=0.77±0.09 and IoU=0.64±0.11 over the 198 MCF-7 tomograms of the independent external test set. As expected, performance metrics in the 3D case are lower than in the 2D case, as an additional dimension introduces further sources of error. Nonetheless, the attained IoU values can still be considered as very good scores for a 3D segmentation problem, as confirmed by Fig. S2, in which we show that small translations of a 3D object cause big reductions in the IoU values. Importantly, also in the 3D case, only a weak decrease can be observed when comparing the training, validation, internal test, independent internal test, and independent external test (Fig. 5e,f), thus confirming the generalization ability of the proposed Holotomography-driven staining also in a 3D scenario.

### Holotomography-driven staining preserves cellular biophysical properties

Among the various reasons that have seen QPI established as an innovative and consistent microscope technique for the single-cell analysis, its quantitative nature stands out as the main advantage [9,11]. QPI allows indeed to quantify biophysical properties related not only to the cell’s morphology, but also to optical features based on its RI distribution, which is a distinctive fingerprint intrinsically connected to the cell biology [45]. For this reason, while the F1 score and the IoU are objective metrics universally employed to assess the performance of a segmentation network, in QPI it is equally important to assess the ability of the network in preserving the biophysical measurements.

To this aim, for each target QPM and corresponding predicted QPM in the 2D Holotomography-driven staining, we measured the optical properties of the nucleus, its size, and its shape, i.e. respectively, the mean phase value, the 2D equivalent diameter, and the circularity, as described in Methods section. A direct comparison between the distributions of these three features in the target and predicted cases is shown in the violin plots of Fig. 6a-c, which were almost symmetrical in all the considered datasets, visually suggesting that the trained Holotomography-driven CNN was able to replicate with high accuracy the QPI measurements in terms of nuclear phase values, size, and shape. To further confirm this, in Fig. 6a-c we also report the mean absolute percentage error (MAPE) with respect to the ground truth (defined in Methods section), which reached its highest values in the independent internal and external test, as expected. Importantly, MAPEs of the independent internal and external test sets were of the same order of magnitude as the training, validation, and internal test sets, confirming the great generalization ability of the trained Holotomography-driven CNN.

**Fig. 6.**
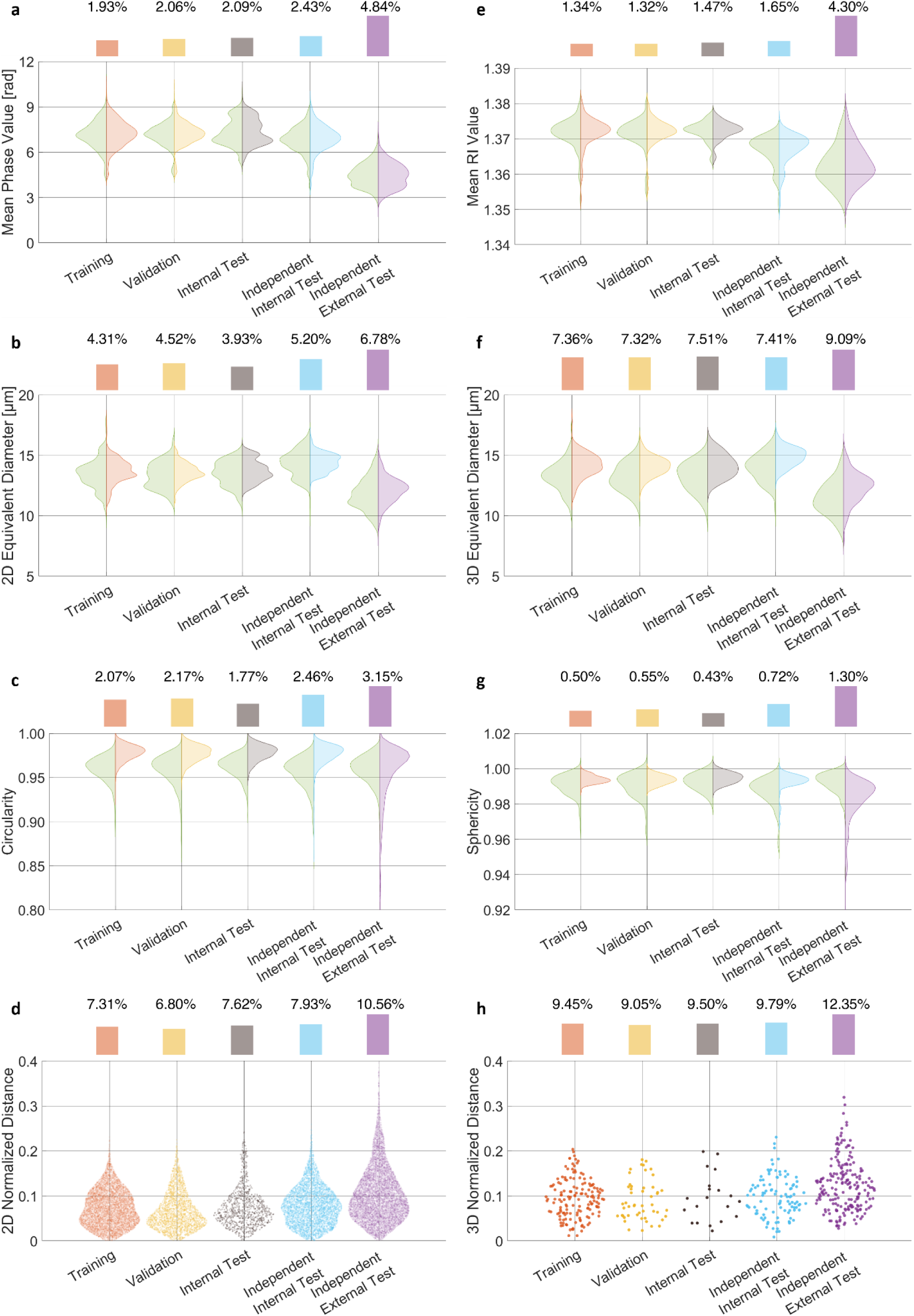
Biophysical features measured in the (a-d) 2D and (e-h) 3D nucleus masks. **a-c,e-g** Violin plots to compare, for each dataset, the ground truth measurements (green) obtained by the CSSI algorithm to the predicted ones obtained by the Holotomography-driven staining. MAPE values are reported at the top. **d,h** Normalized distance between the target nucleus masks and the predicted ones for each dataset. The average values are reported at the top in percentages.

Quantitatively, the average MAPEs among the five datasets were remarkably low, i.e. 2.67% for the mean phase value (Fig. 6a), 4.95% for the 2D equivalent diameter (Fig. 6b), and 2.32% for the circularity (Fig. 6c). This means that, on average, the CNN predicted a slightly bigger and rounder nucleus. Another feature to consider was related to the location of the predicted nucleus inside the cell’s QPM with respect to the target one, which we measured in terms of 2D normalized distance between them (described in Methods section), as reported in the swarm chart of Fig. 6d. Comparable to the examples in Fig. 3a-d, the CNN predicted a 2D nucleus mask closer to the cell’s centroid than the target nucleus mask. This is reflected in a non-negligible 2D normalized distance, which was on average of 8.04%, thus representing the feature with the biggest discrepancy with respect to the ground truth.

We next extended the employed QPI features to the 3D space for assessing the reliability of quantitative biophysical analysis enabled by the 3D Holotomography-driven staining, as reported in Methods section. For each nucleus, we measured the mean RI value (Fig. 6e), its 3D equivalent diameter (Fig. 6f), and its sphericity (Fig. 6g), which differed from the target ones by an average MAPE of 2.02%, 7.74%, and 0.70%, respectively. Moreover, we measured the 3D normalized distance between the 3D nucleus masks of the target and predicted cases to quantify the localization error (described in Methods section), which was on average 10.03%, as reported in Fig. 6h. By comparing the 2D and 3D in-silico staining results, the prediction error was comparable for the mean phase value and the mean RI. Instead, it was lower in terms of sphericity than circularity, while it increased for both the equivalent diameter and the normalized distance in the 3D case than the 2D case.

Finally, in order to carry out a critical evaluation about the realiability and performances of the proposed Holotomography-driven staining, we performed a statistical comparison between the in-silico stained QPMs provided by our HTFC system and the 2D FM images provided by a commercial FIFC system (described in Methods section) of both HeLa cells (Fig. S3) and MCF-7 cells (Fig. S4), as reported in Supplementary section S1. In general, the FM-based assesment confirmed that the Holotomography-driven staining was in good agreement with the in vitro FIFC measurements not only in HeLa model, but also for MCF-7 cells which were never seen during CNN training. In particular, some prediction errors were committed by the Holotomography-driven CNN in terms of nucleus shape and position, but it demonstrated reliable performance in terms of nucleus size.

## Discussion

Bridging the gap in intracellular specificity between QPI and FM is the missing piece needed to establish label-free QPI as a gold-standard tool for advanced single-cell analysis, especially when integrated with flow cytometry. Indeed, technological research in biomedicine is moving fast toward label-free microscopy [7] and deep biophysical cytometry [46], and QPIFC has the potential to be the main character in this process [47]. QPIFC can collect with high-speed thousands of stain-free images of single cells flowing in suspension along a microfluidic channel, by providing at the same time the possibility to extract biophysical features able to phenotype the single-cell fingerprint [45,47]. However, to make the single-cell fingerprint as distinctive as possible, it is crucial to define a strategy for retrieving the missing specificity about the intracellular organelles so that a deep specific intracellular analysis can be carried out. Giant steps in this direction have been made thanks to the revolution introduced by virtual staining based on deep learning [28–35] in the context of in-silico staining [25,26]. However, so far virtual staining has not yet been demonstrated in the flow cytometry environment. Indeed, collecting a dataset of co-registered fluorescent/stain-free images is not trivial in flow cytometry conditions. More importantly, retrieving specificity in a quasi-spherical cell in suspension is more challenging because the intracellular RI contrast is much lower than a cell squeezed at rest on a substrate, in which the cytoplasm is redistributed thus making the intracellular organelles more easily visible even in the absence of staining [27].

Here we have demonstrated that deep learning can be employed to recover intracellular specificity in QPI also in flow cytometry conditions by avoiding any experimental co-registration with fluorescence maps. To do this, we have trained a Holotomography-driven CNN for nucleus segmentation by collecting and using for the first time a fully stain-free dataset of QPMs. Our Holotomography-driven staining has also allowed us to overcome an intrinsic drawback of virtual staining. In fact, virtual staining learns to stain a label-free image from fluorescent images, thus inherently bringing the FM artifacts along related to the staining process itself. Instead, the proposed Holotomography-driven staining is inspired by the framework of self-supervised learning, which is a machine learning approach where labels are generated automatically from the data itself [39], thus avoiding any bias related to exogeneous labelling such as that of medical experts or fluorescent tags. Indeed, labels are typically generated by the model itself or by using algorithms exploiting the inherent structure or relationships within the input data. Therefore, self-supervised learning is making its way in medicine and healthcare [40], with some initial applications also in QPI [41] and static HT [42]. In our case study, to train the Holotomography-driven CNN, we have exploited our HTFC system to create the stain-free dataset of nucleus-specific QPMs. Starting from the 3D RI tomograms of flowing cells, we have computed the stain-free nucleus by the CSSI algorithm, and we have reprojected it along the same viewing angles of the recorded QPMs (Fig. 2a). As summarized in Table S2, the Holotomography-driven CNN, trained on a dataset of HeLa QPMs, has reached an F1 score of 0.88±0.05 and an IoU of 0.80±0.08 on the internal test set of HeLa QPMs. Moreover, a very good generalization has been obtained, even with a second cell line never seen during training. In fact, the Holotomography-driven CNN has achieved an F1 score of 0.87±0.05 and an IoU of 0.78±0.08 on the independent internal test set of HeLa QPMs, and an F1 score of 0.84±0.07 and an IoU of 0.73±0.10 on the independent external test set of MCF-7 QPMs (Table S2).

Remarkably, following the HTFC working principle, by combining the in-silico stained QPMs of a flowing and rolling cell, its 3D RI tomogram can be reconstructed with a retrieved 3D nucleus specificity, thus avoiding the CSSI step (Fig. 4). With reference to the 3D case, the Holotomography-driven CNN has achieved an F1 score of 0.82±0.07 and an IoU of 0.70±0.11 on the internal test set of HeLa tomograms, an F1 score of 0.81±0.07 and an IoU of 0.68±0.09 on the independent internal test set of HeLa tomograms, and an F1 score of 0.77±0.09 and an IoU of 0.64±0.11 on the independent external test set of MCF-7 tomograms (Table S2). It is expected that segmentation metrics in the 3D case are slightly worse than the 2D case. Nevertheless, beyond the pure segmentation metrics, the Holotomography-driven staining has been able to preserve with good accuracy the biophysical features of single flowing cells measured from their QPMs or 3D RI tomograms (Fig. 6), which is a fundamental aspect toward a deep biophysical cytometry. In particular, by considering the biophysical features in Fig. 6, it can be seen that the main errors are partially related to the shape retrieval and mainly related to the localization of the nucleus made by the Holotomography-driven CNN, which cause a decrease in the segmentation metrics (i.e. F1 score and IoU) as a consequence, as also demonstrated in Fig. S1 and Fig. S2. Moreover, even if the 2D QPM measurements obtained by Holotomography-driven staining have been demonstrated to be in good agreement with the 2D FM measurements obtained by a commercial FIFC system, the FM assessment in Fig. S3 and Fig. S4 has confirmed the same error sources found by the QPI assessment in Fig. 6.

It is worth remarking that the in-silico staining based on Holotomography-driven learning can overcome the main limitation of the CSSI algorithm, i.e. its long and heavy computations. Indeed, the proposed CNN, derived from a DeepLabv3+ ResNet-18, predicts the 2D nucleus mask in just 2.7 ms after taking in input the QPM, meaning that the 3D nucleus mask can be computed in the order of tenths of a second instead of tens of minutes. Furthermore, the employed CNN has an extremely lightweight architecture, as reported in Methods section, with only 29.6k learnable parameters corresponding to 128 KB of memory occupation. This means that the proposed strategy can go into the direction of real-time and on chip in-silico staining of QPMs and 3D RI tomograms of stain-free cells in flow cytometry environment. Of course, better segmentation performance can be obtained by creating and training ad-hoc CNN architectures for this specific problem. As we believe the proposed in-silico staining based on Holotomography-driven learning could represent in the future a decisive step towards the ambitious deep biophysical cytometry, we have shared with the scientific community the stain-free dataset of QPMs to support a faster advancement in this field.

## Methods

### Sample preparation

MCF-7 (human breast cancer cells) and HeLa (cervical cancer) cells were cultured in Dulbecco’s Modified Eagle’s Medium (DMEM, Gibco, #11995-065) supplemented with 20% and 10% fetal bovine serum (FBS, Euroclone, #ECS0186L), respectively, along with penicillin (100 IU/mL) and streptomycin (100 μg/mL) (Euroclone, #ECB3001D). For cell harvesting, cultures were incubated with 0.05% trypsin-EDTA solution (Sigma-Aldrich) for 5 minutes to promote detachment. Cells were then resuspended in complete culture medium and adjusted to a final concentration of 3 × 10⁵ cells/mL. Subsequently, the cell suspension was introduced into a microfluidic channel for HTFC experiments.

For FM analysis, 3 million HeLa and MCF-7 cells were washed and resuspended in 1× phosphate-buffered saline (PBS) (Sigma) followed by nuclei staining with 25 μM DRAQ5 fluorescent probe (#62254, Thermo Scientific™) for 5 minutes at room temperature under agitation.

### Holo-Tomographic Flow Cytometry: experimental system and numerical processing

To realize the HTFC experimental setup, we implemented a Mach-Zehnder interferometer based on off-axis configuration [23], as displayed in Fig. S6a. The light wave, generated by a solid state continuous wave laser (Laser Quantum Torus - 532, λ = 532 nm), was split by a polarizing beam splitter (PBS) in reference and objective beams. The reference beam was transmitted, while the objective beam was reflected. The objective beam passed through the cells flowing and rotating along the microfluidic channel (MC; Microfluidic ChipShop). Hence, the scattered light was collected by a microscope objective (MO1; Zeiss, ×40, oil immersion, NA = 1.3) and was driven, by a mirror (M4), to a tube lens (TL1, focal distance = 150 mm). The refence beam was driven through a second microscope objective (MO2) and a second tube lens (TL2), which acted as a beam expander. According to the definition of off-axis configuration, the two contributions were recombined by a beam splitter (BS) with a non-null angle. The resulting interference fringe pattern was collected by a conventional CMOS (Genie Nano-CXP Camera, 5120 × 5120 pixels, Δx = Δy = 4.5 μm pixel size). Holographic videos were recorded with a frame rate of 30 frame per second (fps). As the overall lateral magnification in the camera plane reached 36, the recorded field of view (FoV) measured 640 × 640 μm^2^, as shown in Fig. S6b. In order to control the flow, we used an automatic syringe pump (CETONI Syringe Pump neMESYS 290N) for injecting cells into the MC. Within the channel, a laminar flow was generated and a quasi-uniform flow was guaranteed. Cells not flowing in the center of the MC underwent the velocity gradient due to the parabolic velocity profile, thus starting to rotate. Given the Cartesian reference system in Fig. 6b, cells flowed along the y-axis, rotated around the x-axis, and were imaged along the z-axis.

The acquired holographic videos were used to track each flowing cell (Fig. S6b). In this way, for each flowing and rotating cell, a stack of sub-holograms (Fig. S6c) was available at different rolling angles (i.e., viewing angles). To reconstruct the QPM corresponding to each sub-hologram [23], we first demodulated it by isolating the desired +1 diffraction order through a band-pass filter in the Fourier domain. Then, to achieve optimal focus, we minimized the Tamura coefficient along the optical axis, and we refocused the complex field by numerically propagating the demodulated sub-hologram through the Angular Spectrum method. Hence, we performed aberration compensation by subtracting a reference hologram (i.e. a hologram without cells inside), phase denoising through the 2D windowed Fourier transform, and phase unwrapping through the PUMA algorithm, thus obtaining the corresponding QPM (Fig. S6d). For each flowing and rotating cell, the rolling angles related to each QPMs were estimated by a phase-contrast image similarity metric, and finally the 3D RI tomograms were reconstructed through the FBP algorithm [23].

### Fluorescence Imaging Flow Cytometry

To record thousands of 2D FM images, we employed a commercial multispectral FIFC system, specifically the ImageStream^®^X Mark II flow cytometer (Luminex Corporation). Using this system, we acquired around 1,000 single-cell images at 40× magnification for each cell line. Data acquisition was performed using the INSPIRE software. Cells were identified based on brightfield images (Fig. S3a and Fig. S4a) and nuclear signals (Fig. S3b and Fig. S4b) using a 642 nm red laser (20 mW) to detect the DRAQ5 fluorescence dye. After analysing the single-cell images using IDEAS software (version 6.2.64.0), we deleted false positive, thus obtaining 362 HeLa cells and 231 MCF-7 cells. Finally, a double threshold computed by the Otsu’s method from the fluorescence images allowed to segment both the cell contour and the nucleus contour (Fig. S3c and Fig. S4c), which were then employed to assess the proposed in-silico staining strategy.

### CNN training and performance evaluation

The MATLAB^®^ R2024b software was employed to train the CNN for the in-silico staining. The CNN model was derived from the built-in DeepLabv3+ ResNet-18 architecture (*deeplabv3plus* function) [43]. However, it was adapted for the in-silico staining problem by performing some changes. First, the input size was set to 96×96×1 to take in input the QPMs provided by the HIFC system (Fig. 3a-d), and the input QPMs were normalized in the [0,1] range to help generalization. Moreover, the architecture was lightened by removing the 4 convolutional layers before the atrous spatial pyramid pooling (ASPP), namely *res4a*, *res4b*, *res5a*, and *res5b*. During training, the CNN used as target a binary image in which the nucleus mask was set to 1 (Fig. 3a-d), and tried to minimize a loss function made of the sum of three terms, i.e.

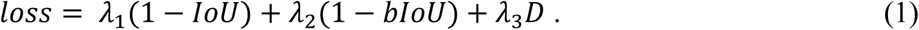

The first loss term is based on the IoU [48], also known as Jaccard index, defined as the ratio of the intersection and union areas between the target and predicted nucleus masks, i.e.

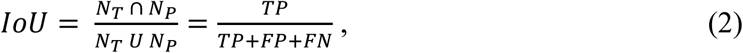

where *N_T_* is the area of the target nucleus mask, *N*_*P*_ is the area of the predicted nucleus mask, *TP* is the number of nucleus pixels correctly segmented as nucleus by the CNN (i.e. true positives), *FP* is the number of non-nucleus pixels wrongly segmented as nucleus by the CNN (i.e. false positives), and *FN* is the number of nucleus pixels wrongly segmented as non-nucleus by the CNN (i.e. false negatives). The IoU is 1 if there is a perfect overlapping between the target and predicted nucleus mask, while it is 0 if there is no overlapping between the target and predicted nucleus mask. Hence, the IoU loss term allowed to train the CNN to segment well the central area of the nucleus mask.

The second loss term is based on the boundary IoU (bIoU) [49], which was employed to train the CNN to segment well the contours of the nucleus mask. To compute the boundary IoU, a boundary nucleus mask was defined by first isolating the nucleus contour through a 3×3 Laplacian filter, and then by dilating it through a 5×5 average filter. Finally, the IoU was computed over the boundary regions of the target and predicted nucleus masks.

The third loss term is the Euclidean distance *D* between the centroids target and predicted nucleus mask, and it was employed to train the CNN to localize well the nucleus mask.

Furthermore, to help generalization, at each iteration of the training, a random spatial translation was added to the input and target images of the training set.

The CNN was trained for 100 epochs by using a Desktop computer mounting an Intel^®^ Core™ i9-9900K CPU @ 3.60 GHz, a 64 GB RAM, and an NVIDIA Quadro P1000 GPU. During training, 10^−4^ was used as learning rate, 256 as minibatch size, 5 as validation frequency, 0.1 as *λ*_1_ and *λ*_2_, and 0.8 as *λ*_3_. The lightweight CNN had a reduced number of learnable parameters (29.6k) with respect to the original ResNet-18 (20.6M), thus allowing to optimize the training time (∼2.5 h), the memory occupation (128 KB), and the prediction time (2.7 ms). The training and validation loss are shown in Fig. S5. The best validation loss was used as a criterion to select the trained model throughout the 100 epochs.

To evaluate the performance of the segmentation network, the IoU and the F1 score were used as metrics. The F1 score, also known as Dice coefficient, is the harmonic mean between recall and precision [50], i.e.

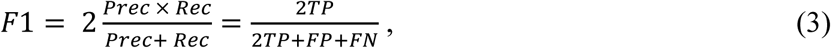

where the recall (Rec) is the probability of correctly segmenting a pixel belonging to the nucleus, and the precision (Prec) is the probability that a pixel predicted as nucleus is correct. It is worth noting that the nucleus mask predicted by the trained CNN was convolved with a 7×7 average kernel, in order to delete residual errors and smooth the boundaries. This operation was performed before evaluating the performance metrics.

### Feature analysis

The MATLAB^®^ R2024b software was employed to measure the morphological and biophysical features from the 2D and 3D nucleus masks discussed in Fig. S3, Fig. S4, and Fig. 6.

In particular, to compare the FM measurements to the in-silico staining ones in the 2D case (Fig. S3 and Fig. S4), three features were computed, i.e. the nucleus-cell area ratio, the nucleus circularity, and the nucleus-cell centroids normalized distance. The nucleus-cell area ratio is the ratio between the nucleus area and the cell area in a 2D image. The nucleus circularity is defined as

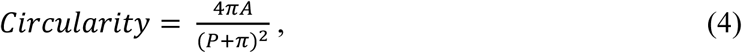

where *A* and *P* are the area and perimeter of the 2D nucleus mask. Circularity is the ratio of the perimeter of a circle having the same area of the 2D nucleus mask to the 2D nucleus mask’s perimeter. Thus, circularity is 1 for a perfect circle, otherwise it is lower than 1. The nucleus-cell centroids normalized distance is the distance between the nucleus centroid and the cell centroid in a 2D image, normalized to the 2D equivalent diameter of the cell. In general, the 2D equivalent diameter of a 2D object is the diameter of a circle having the same area as the 2D object mask. To compute the dispersion ellipsoid related to these three features (Fig. S3g and Fig. S4g), they were first standardized by subtracting their median and dividing by their interquartile range. Then, for each feature, the mean and standard deviation values were computer, which were finally employed as centres and semiaxes of the dispersion ellipsoid, respectively.

To compare the CSSI measurements to the Holotomography-driven learning ones in the 2D and 3D cases (Fig. 6), six nuclear features were computed, i.e. the mean phase value, the mean RI value, the 2D equivalent diameter, the 3D equivalent diameter, the circularity, and the sphericity. The mean phase value is the average QPM value computed inside the 2D nucleus mask. The mean RI value is the average RI value computed inside the 3D nucleus mask. The 3D equivalent diameter is the diameter of a sphere having the same volume as the 3D nucleus mask. The sphericity is defined as

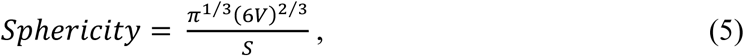

where *V* and *S* are the volume and the surface area of the 3D nucleus mask. Sphericity is the ratio of the surface area of a sphere having the same volume of the 3D nucleus mask to the 3D nucleus mask’s surface area. Thus, sphericity is 1 for a perfect sphere, otherwise it is lower than 1. For these six features, the prediction error was computed by MAPE, defined as

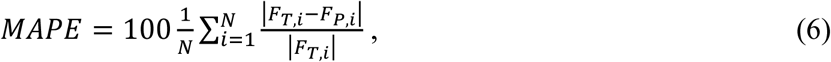

where *F*_*T*_ and *F*_*P*_ are the feature values computed from the target and predicted nucleus masks, respectively, and *N* is the number of observations.

Finally, the 2D and 3D normalized distances are discussed in Fig. 6. The 2D normalized distance is the distance between the centroids of the target and predicted 2D nucleus masks, normalized to the 2D equivalent diameter of the target 2D nucleus mask. Instead, the 3D normalized distance is the distance between the centroids of the target and predicted 3D nucleus masks, normalized to the 3D equivalent diameter of the target 3D nucleus mask. In both cases, the prediction error was computed as their average value expressed in percentages.

## Data availability

The dataset of 2D QPMs employed to train and test the Holotomography-driven CNN is publicly available in the GitHub repository https://github.com/danpir94/Holotomography_driven_learning_for_in_silico_staining.

## Acknowledgements

This work was supported by project PRIN 2022 PNRR – Flow-cytometry ImaGing by Holographic tomography for predicting TUMor control in Oncology patients treated with Radiotheraphy (FIGHT-TUMOR), Prot. P2022ATE2J – funded by the Italian Ministry of University & Research in the framework of European Union program Next Generation EU (CUP: B53D23023890001), and by project PRIN 2022 – Computationally aided Opto-mechano-fluidic pLatform for Label-free intelligent tumor microEnvironment Cell sorTing (COLLECT) Prot. 202275PJRP – funded by the Italian Ministry of University & Research in the framework of the European Union program Next Generation EU (CUP: B53D23002280006).

## Author contributions

D.P., P.M., and P.F. conceptualized the idea behind the study and wrote the original draft. L.M. and P.M. designed the holographic experiments and the holographic processing, respectively. M.S., A.M., and M.M. conducted cell cultures. G.G. performed holographic experiments. V.C. and M.R. performed cytofluorimetric experiments. D.P. designed the computational methods behind the study, conducted formal analysis and validated results. A.M., I.K., D.L.M., L.M., and M.C. revised the manuscript. I.K., D.L.M., M.C., and A.I. supervised the biological experiments. G.S. supervised the cytofluorimetric experiments. L.M. supervised the holographic experiments. P.M. and P.F. supervised the study and managed the acquisition of funding. All authors red and approved the final version of the manuscript.

## Competing interests

The authors declare no competing interests.

## Supplementary information

**Supplementary Video 1 –** Workflow of the Holotomography-driven learning for the in-silico staining of 2D QPMs and 3D RI tomograms of single cells flowing along a microfluidic channel avoiding co-registration (MP4).

## Notes

https://github.com/danpir94/Holotomography_driven_learning_for_in_silico_staining

### Competing Interest Statement

The authors have declared no competing interest.

