## Supplementary Information for "Holotomography-driven learning for in-silico staining of single cells in flow cytometry avoiding co-registration"

### S1. Comparison between FM imaging and Holotomography-driven staining.

We collected two FIFC datasets, one formed by 362 HeLa cells and the other one formed by 231 MCF-7 cells, and we respectively compared them to the QPIFC datasets of 323 HeLa cells (i.e., the HeLa cells of the training, validation, internal test, and independent internal test sets, Table S1) and 198 MCF-7 cells (i.e., the MCF-7 cells of the independent external test set, Table S1). It is important to note that, for a fair comparison, to create the QPIFC dataset, for each flowing and rolling cell recorded by the HTFC system (Table S1), we considered only the first QPM of the entire stack. Moreover, as for the FIFC dataset, for each cell, we recorded the brightfield image of the cell (Fig. S3a and Fig. S4a) and the DRAQ5 image of the nucleus (Fig. S3b and Fig. S4b), thus obtaining the segmented cell displayed in Fig. S3c and Fig. S4c. Instead, to segment the stain-free QPMs, we exploited both the CSSI algorithm and the proposed Holotomography-driven CNN.

From both the FM images and the QPMs, we measured the nucleus size, shape, and position in terms of nucleus-cell area ratio, nucleus circularity, and nucleus-cell centroids normalized distance, respectively, as described in Methods section. For each feature, to perform a statistical comparison, we implemented the Wilcoxon-Mann-Whytney (WMW) non-parametric statistical hypothesis test [S1], as shown in the boxplots of Fig. S3d-f and Fig. S4d-f. As expected, for all three features, the CSSI algorithm showed non-significant p-values (i.e., higher than the significance level of 0.01). Remarkably, the nucleus size exhibited a non-significant p-value also with the Holotomography-driven staining in both HeLa (Fig. S3d) and MCF-7 (Fig. S4d) cells. As for the shape, the in-silico stained nucleus appeared more circular than the FM one in the HeLa case (p-value≤0.00001, Fig. S3e), while the nucleus circularity of the in-silico stained nucleus was on average lower than the FM one in the MCF-7 case (p-value≤0.001, Fig. S4e). Furthermore, the in-silico stained nucleus was closer to the cell centroid than the FM one in the HeLa case (p-value≤0.00001, Fig. S3f), while the in-silico stained nucleus was almost in the same position as the FM one in the MCF-7 case (non-significant p-value, Fig. S4f).

Finally, to perform an overall comparison in terms of nucleus size, shape, and position with respect to the gold-standard FIFC, we built the dispersion ellipsoids starting from the marginal distributions of the three measured features (as reported in Methods section), as shown in Fig. S3g and Fig. S4g. The dispersion ellipsoid of the CSSI algorithm, which was in strong agreement with the FIFC measurements according to the p-value analysis, reached an IoU of 0.68 in the HeLa cells (Fig. S3g) and 0.67 in the MCF-7 cells (Fig. S4g) with respect to the FM dispersion ellipsoid. Instead, the dispersion ellipsoid of the Holotomography-driven staining reached an IoU of 0.61 in the HeLa cells (Fig. S3g) and 0.59 in the MCF-7 cells (Fig. S4g) with respect to the FM dispersion ellipsoid.

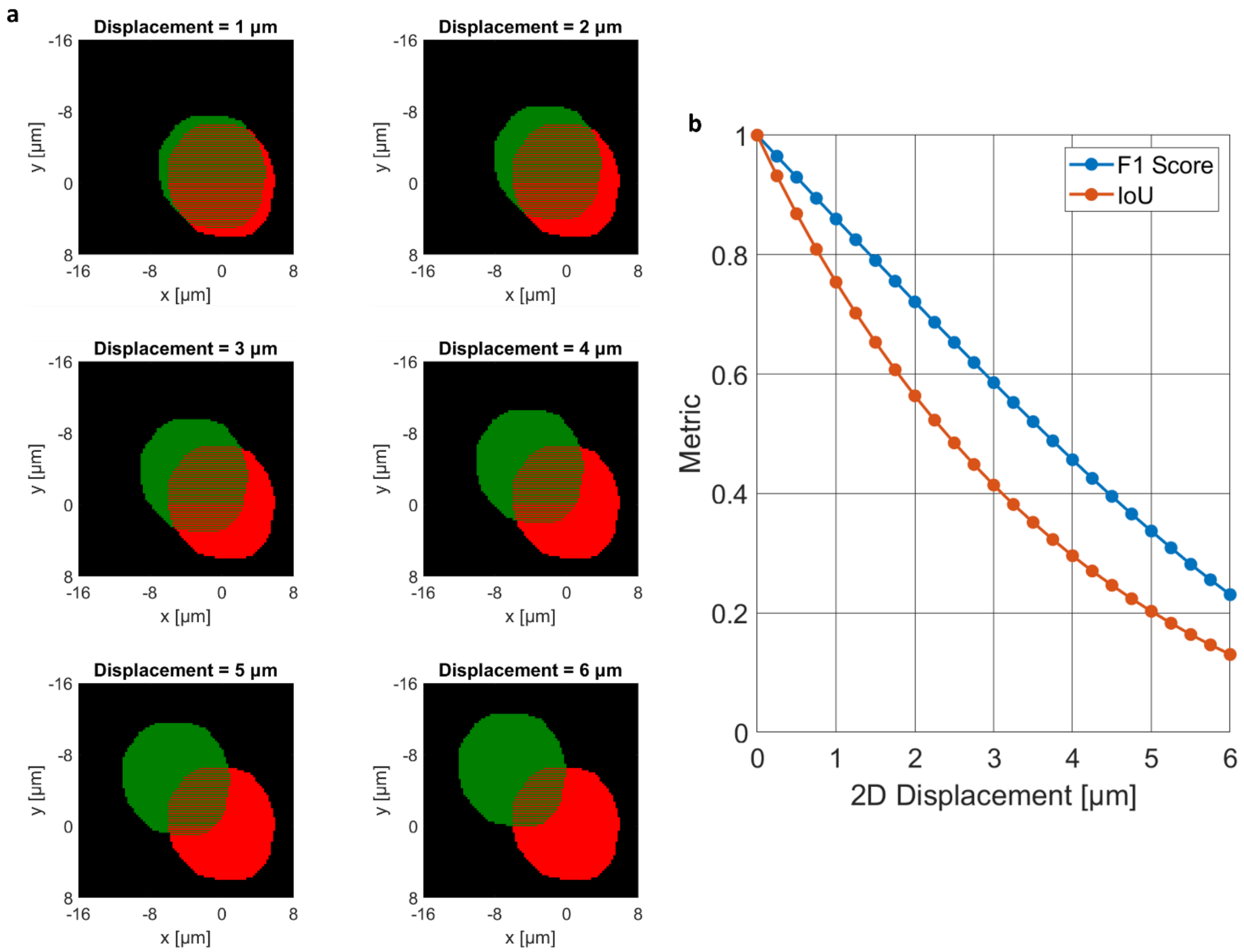


**Fig. S1. Effect on the F1 score and IoU metrics of the sole displacement of a 2D nucleus mask. a** Examples of displacement. The original 2D nucleus mask (red) is displaced in the x-y directions by the values reported at the top, thus obtaining the translated 2D nucleus mask (green). **b** F1 score and IoU metrics at different displacements.

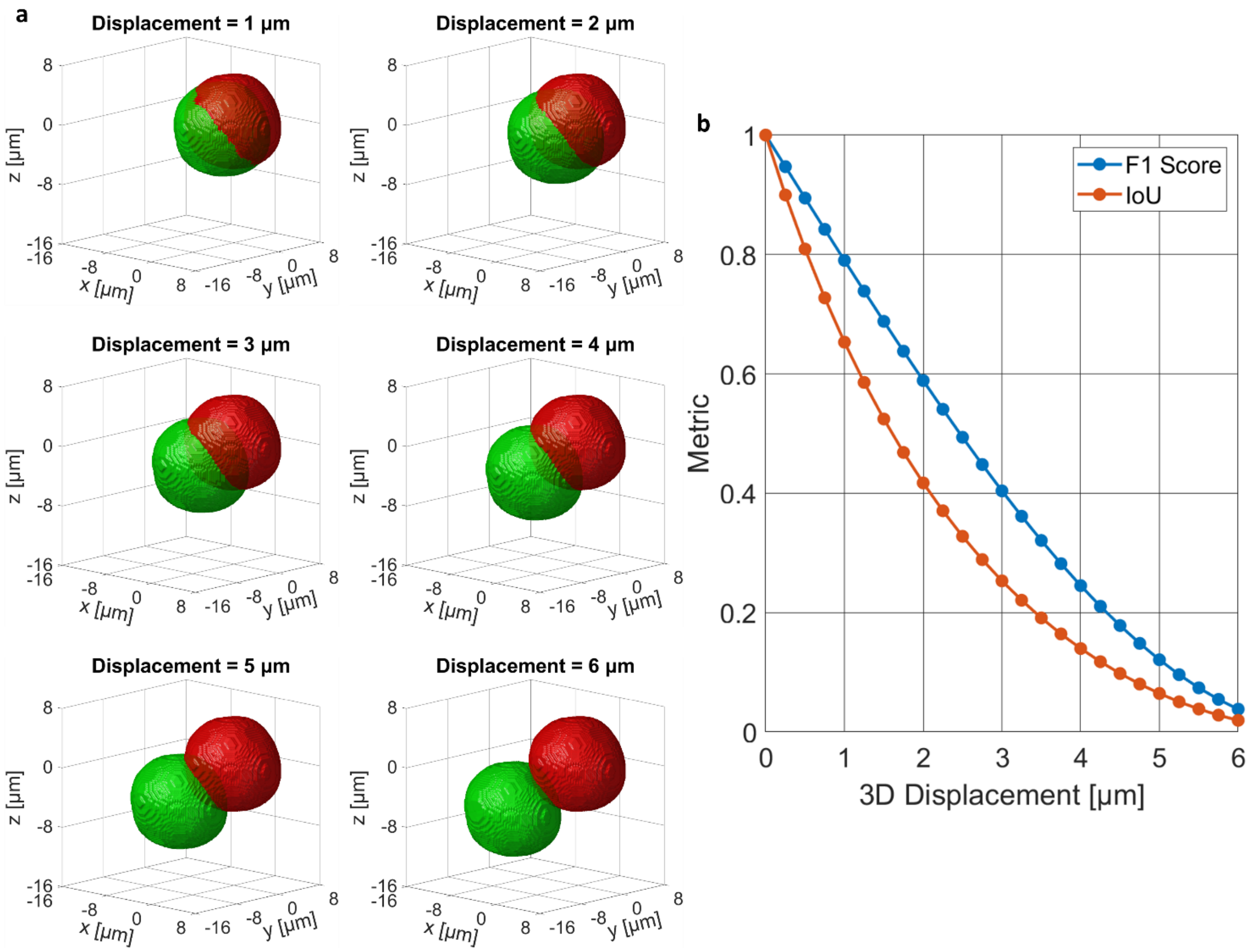


**Fig. S2. Effect on the F1 score and IoU metrics of the sole displacement of a 3D nucleus mask. a** Examples of displacement. The original 3D nucleus mask (red) is displaced in the x-y-z directions by the values reported at the top, thus obtaining the translated 3D nucleus mask (green). **b** F1 score and IoU metrics at different displacements.

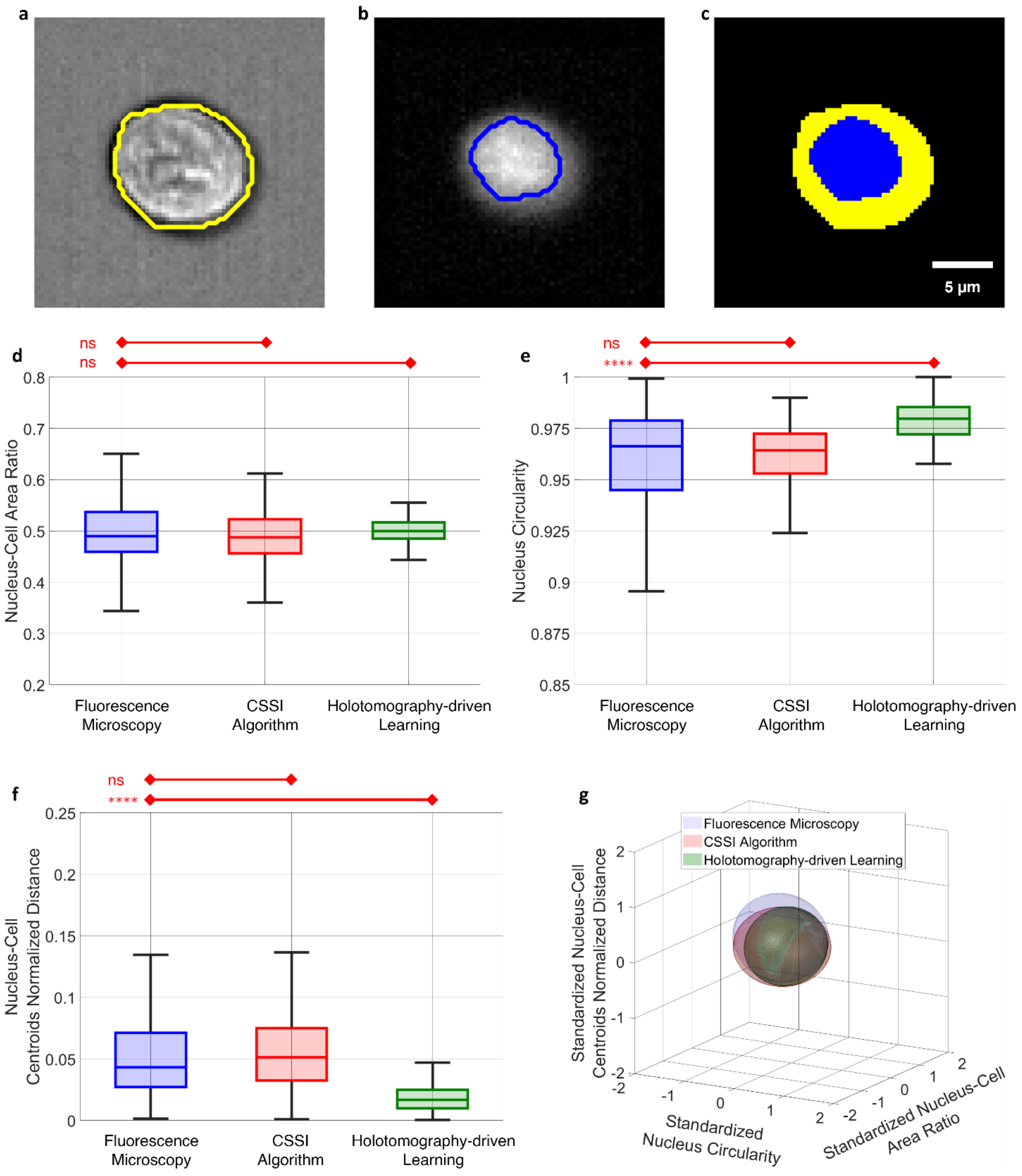


**Fig. S3. FM assessment of the in-silico staining of the nucleus in HeLa cells based on FIFC measurements.** **a** Example of brightfield image obtained by FIFC, with highlighted the segmented cell contour (yellow). **b** Example of DRAQ5 image obtained by FIFC, with highlighted the segmented nucleus contour (blue). **c** Nucleus binary mask (blue) in (b) overlapped to the cell binary mask (yellow) in (a). **d-f** Boxplots of the nucleus-cell area ratio, nucleus circularity, and nucleus-cell centroids normalized distance, respectively, to compare the FM measurements (n=362) to the in-silico staining measurements (n=323) obtained by the CSSI algorithm and the Holotomography-driven learning in terms of nucleus size, shape, and position, respectively. The corresponding p-values are reported at the top. For each boxplot, the central line is the median, the bottom and top box edges are the 25th and 75th percentiles, respectively, and the whiskers extend to the most extreme non-outlier data points. ns-not significant p>0.01, **** p<0.00001. **g** Dispersion ellipsoids of the FM standardized measurements (blue) and the in-silico staining standardized measurements obtained by the CSSI algorithm (red) and the Holotomography-driven learning (green). The IoU between the FM and CSSI dispersion ellipsoids is 0.68, and the IoU between the FM and Holotomography-driven staining dispersion ellipsoids is 0.61.

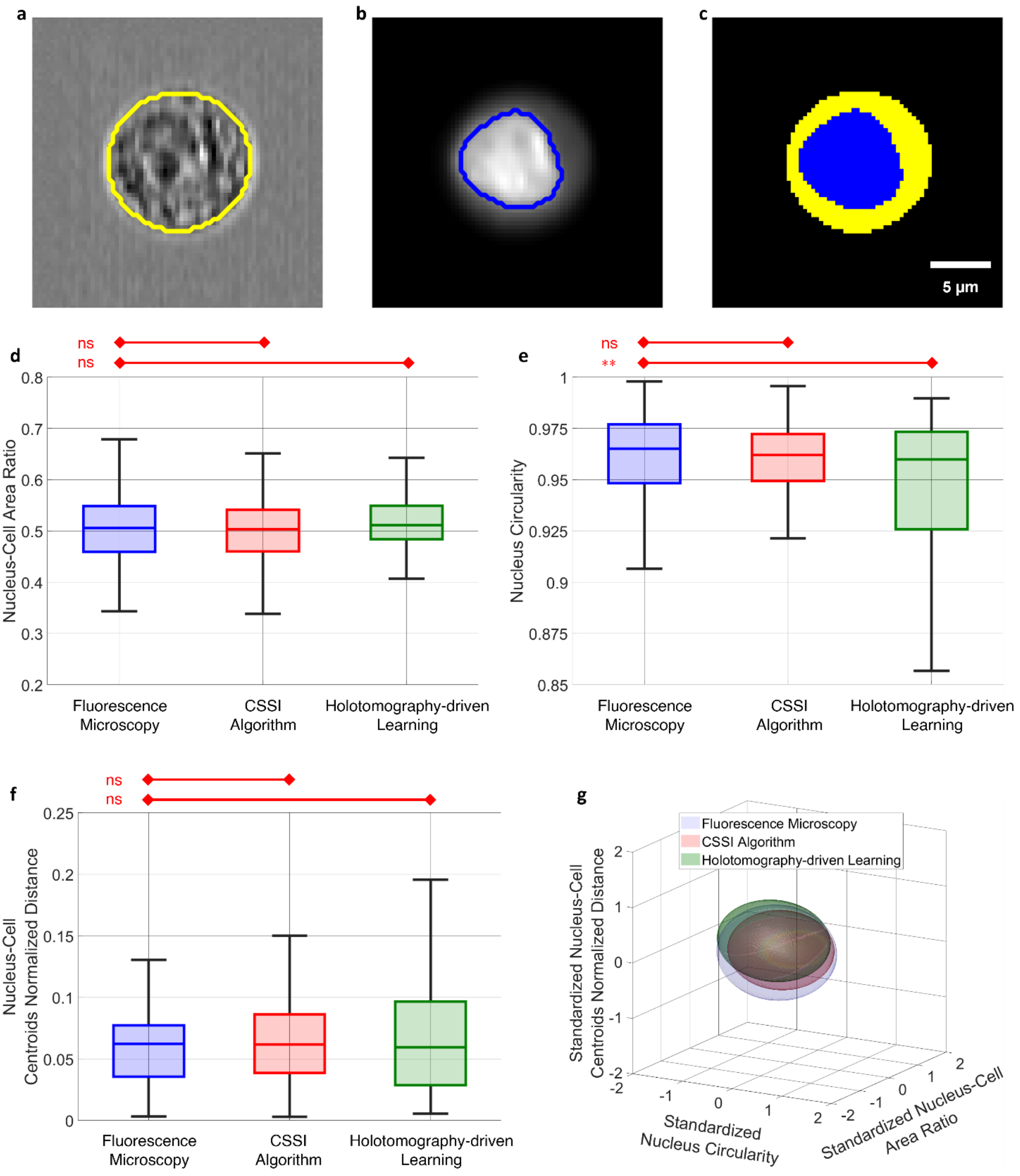


**Fig. S4. FM assessment of the in-silico staining of the nucleus in MCF-7 cells based on FIFC measurements. a** Example of brightfield image obtained by FIFC, with highlighted the segmented cell contour (yellow). **b** Example of DRAQ5 image obtained by FIFC, with highlighted the segmented nucleus contour (blue). **c** Nucleus binary mask (blue) in (b) overlapped to the cell binary mask (yellow) in (a). **d-f** Boxplots of the nucleus-cell area ratio, nucleus circularity, and nucleus-cell centroids normalized distance, respectively, to compare the FM measurements (n=231) to the in-silico staining measurements (n=198) obtained by the CSSI algorithm and the Holotomography-driven learning in terms of nucleus size, shape, and position, respectively. The corresponding p-values are reported at the top. For each boxplot, the central line is the median, the bottom and top box edges are the 25th and 75th percentiles, respectively, and the whiskers extend to the most extreme non-outlier data points. ns-not significant $p>0.01$, **** $p<0.00001$. **g** Dispersion ellipsoids of the FM standardized measurements (blue) and the in-silico staining standardized measurements obtained by the CSSI algorithm (red) and the Holotomography-driven learning (green). The IoU between the FM and CSSI dispersion ellipsoids is 0.67, and the IoU between the FM and Holotomography-driven staining dispersion ellipsoids is 0.59.

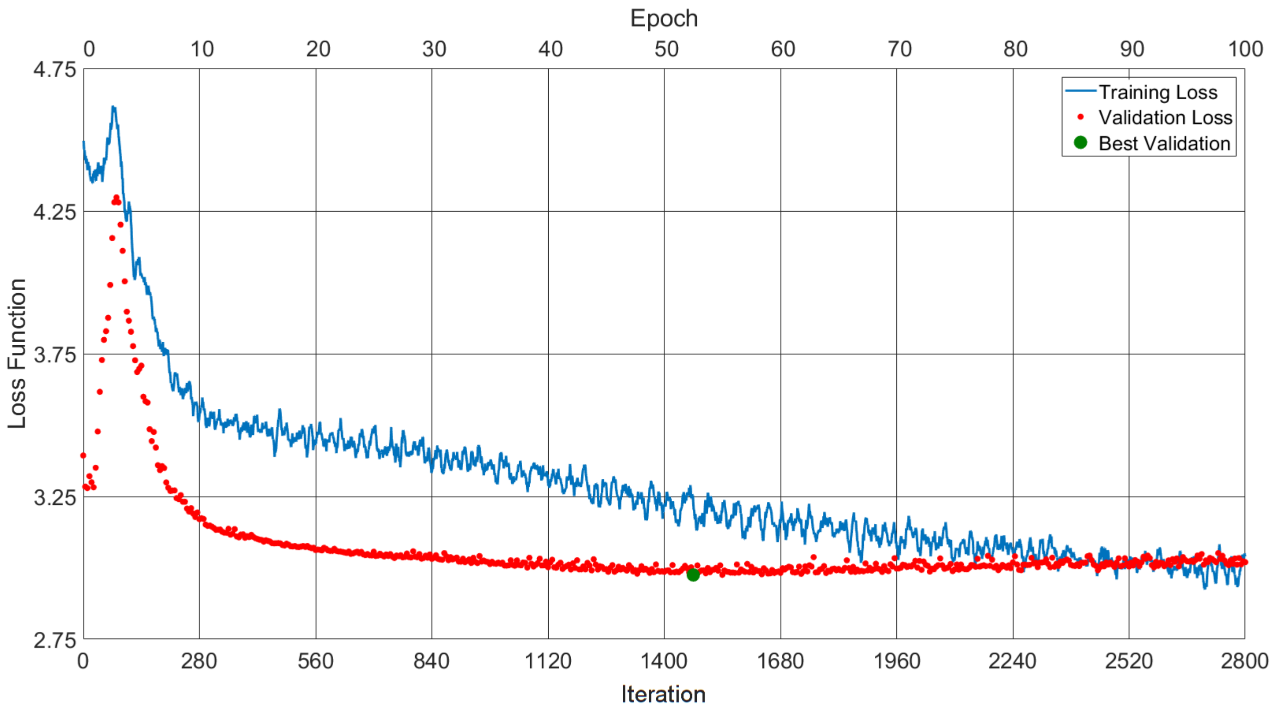


**Fig. S5. Loss function during the training of the in-silico staining CNN.** The best validation loss (green dot) is used as criterion to select the trained model.

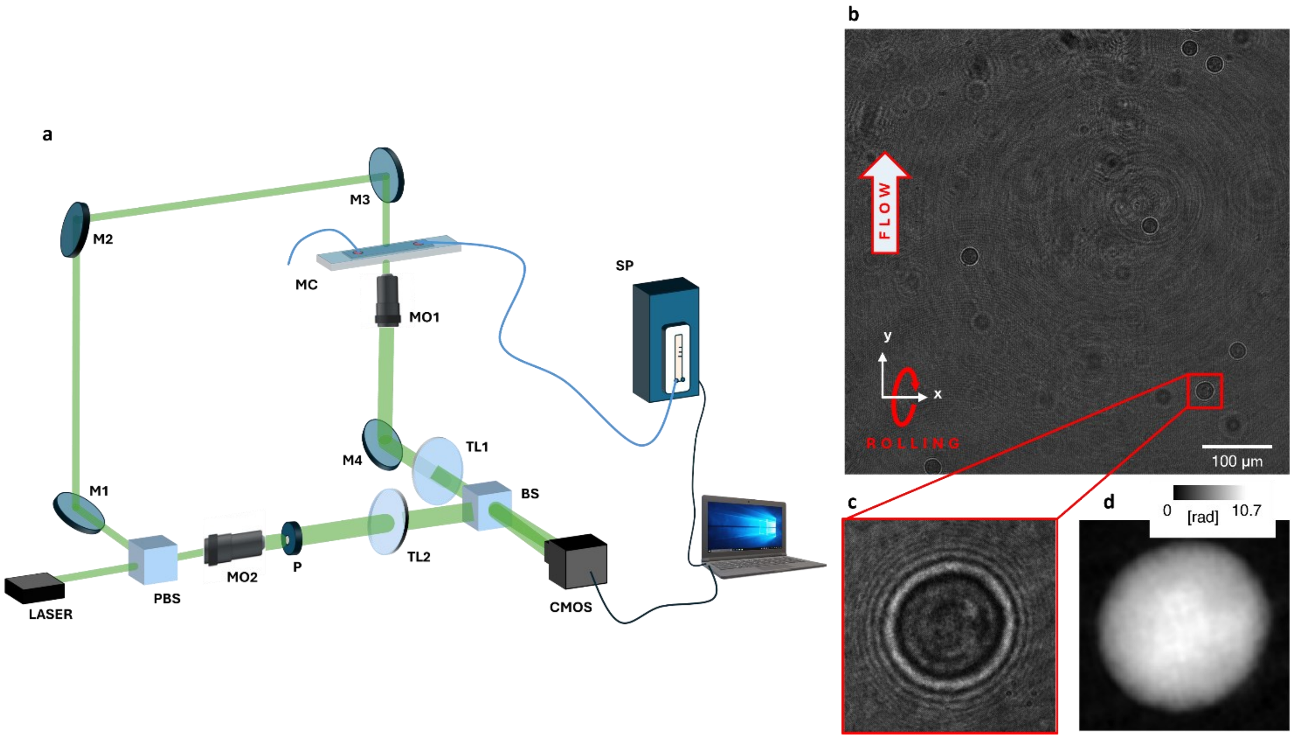


**Fig. S6. HTFC working principle. a** Sketch of the opto-fluidic experimental system. PBS – polarizing beam splitter; M – mirror; MO – microscope objective; P – pinhole; TL – tube lens; BS – beam splitter; MC – microfluidic channel; SP – syringe pump; CMOS – camera. **b** Digital hologram taken from the holographic video sequence recorded by the HTFC system in (a). Cells flow along the y-axis, rotate around the x-axis, and are imaged along the z-axis. **c** Sub-hologram containing the cell highlighted in (b). **d** QPM computed from the sub-hologram in (c).

**Table S1.** Stain-free dataset of single flowing cells collected by HTFC.

| | *Type* | *Cell Line* | *# Cells* | *# QPMs* |
|---|---|---|---|---|
| *Training* | Training Set | HeLa | 150 | 7,297 |
| | Validation Set | HeLa | 50 | 2,418 |
| | Internal Test Set | HeLa | 20 | 1,014 |
| | | | *220* | *10,729* |
| *Test* | Independent Internal Test Set | HeLa | 103 | 4,415 |
| | Independent External Test Set | MCF-7 | 198 | 8,255 |
| | | | *301* | *12,670* |

**Table S2.** Segmentation metrics to evaluate performances of the Holotomography-driven CNN.

| | *Type* | *Cell Line* | *2D QPMs* | | *3D RI Tomograms* | |
|---|---|---|---|---|---|---|
| | | | *F1 Score* | *IoU* | *F1 Score* | *IoU* |
| *Training* | Training Set | HeLa | 0.89 ± 0.04 | 0.80 ± 0.07 | 0.82 ± 0.06 | 0.71 ± 0.09 |
| | Validation Set | HeLa | 0.89 ± 0.05 | 0.80 ± 0.07 | 0.82 ± 0.06 | 0.70 ± 0.09 |
| | Internal Test Set | HeLa | 0.88 ± 0.05 | 0.80 ± 0.08 | 0.82 ± 0.07 | 0.70 ± 0.11 |
| *Test* | Independent Internal Test Set | HeLa | 0.87 ± 0.05 | 0.78 ± 0.08 | 0.81 ± 0.07 | 0.68 ± 0.09 |
| | Independent External Test Set | MCF-7 | 0.84 ± 0.07 | 0.73 ± 0.10 | 0.77 ± 0.09 | 0.64 ± 0.11 |
